# The ecology and quantitative genetics of seed and seedling traits in upland and lowland ecotypes of a perennial grass

**DOI:** 10.1101/2021.12.14.472686

**Authors:** Samsad Razzaque, Thomas E Juenger

## Abstract

Plants have evolved diverse reproductive allocation strategies and seed traits to aid in dispersal, persistence in the seed bank, and establishment. In particular, seed size, dormancy, and early seedling vigor are thought to be key functional traits with important recruitment and fitness consequences across abiotic stress gradients. Selection for favored seed-trait combinations, or against maladaptive combinations, is likely an important driver shaping recruitment strategies. Here, we test for seed-trait plasticity and local adaptation in contrasting upland and lowland ecotypes of *Panicum hallii* with field experiments in native versus foreign habitats. Furthermore, we test whether seed traits have been under directional selection in *P. hallii* using the *v*-test (Fraser 2020) based on trait variance in a genetic cross. Finally, we evaluate the genetic architecture of ecotypic divergence for these traits with Quantitative Trait Locus (QTL) mapping. Field experiments reveal little plasticity but support a hypothesis of local adaptation among ecotypes based on recruitment. Patterns of segregation within recombinant hybrids provides strong support for directional selection driving ecotypic divergence in seeds traits. Genetic mapping revealed a polygenic architecture with evidence of genetic correlation between seed mass, dormancy, and seedling vigor. Our results suggest that the evolution of these traits may involve constraints that affect the direction of adaptive divergence. For example, seed size and germination percentage shared two colocalized QTL with antagonistic additive effects. This supports the hypothesis of a functional genetic relationship between these traits, resulting in either large seed/strong dormancy or small seed/weak dormancy trait combinations. Overall, our study provides insights into the factors facilitating and potentially constraining ecotypic differentiation in seed traits.

**Impact Summary:** Seed size and dormancy are key functional traits with important recruitment and fitness consequences. Theory suggests tradeoffs and plasticity in offspring quantity and quality are important in seed evolution. The genetics of seed size and dormancy traits have been studied extensively, but these studies are mostly limited to model system or domesticated crops. We also know very little about the genetic architecture and evolution of seed-based life history traits especially considering adaptation to xeric and mesic habitats. Here, we explored the genetic basis of trade-offs between seed size and dormancy in a C4 perennial grass, *Panicum hallii*, endemic to North America. We planted seeds of recombinant inbred lines from a cross between a xeric and mesic ecotype of *P*.*hallii* and mapped quantitative trait (QTL) loci for seed size, dormancy and seedling vigor traits. We detected a genetic basis of trade-offs between seed size and dormancy, suggesting that natural selection strongly favored specific trait combinations in ecotype formation. We further explored the role of seed size variation on seedling and adult recruitment in contrasting habitats. Our data showed that seed size was under strong selection through recruitment. Overall, our results demonstrate that adaptive differentiation for seed size and early life stages are important factors in adaptation to contrasting habitats.

## Introduction

Local adaptation is an important component of responses to changing environments, as it reveals how environmental variation can drive phenotypic and genetic differentiation as a response to selection (Davis and Shaw 2001; Conover et al. 2009). In this process, plant populations may be selected for different resource allocation strategies and trait combinations based on habitat structure or climatic features across a species range. One hypothesis is that local adaptation results from genetic trade-offs in which locally favored alleles reduce fitness elsewhere to provide environment specific fitness advantage (Roff 2002). For example, plants have evolved diverse seed and seedling traits to aid adaptation in response to heterogeneity in the amount of rainfall (Smith and Fretwell 1974; Daws et al. 2007; Venable 1992). These trade-offs may maintain genetic variation among populations and drive phenotypic partitions within species (Futuyma and Moreno 1988; Jasmin and Kassen 2007). Thus, exploring the genetic basis of trade-offs and how they shape ecological strategy is therefore of major importance for evolutionary biology (Agrawal et al. 2010; Postma and Ågren 2016).

In plants, soil water availability plays a vital role maintaining global species distribution and determining genetic and phenotypic differences within species range (Stebbins Jr 1952; Whittaker 1970; Woodward et al. 2004). Thus, plant adaptation to habitats differing in soil water availability can drive the evolution of divergence and may promote ecological speciation with distinct morphological and/or physiological characteristics that provide an environment-specific fitness advantage (Baker 1972; Roux et al. 2006; Juenger 2013; Rundle and Nosil 2005). However, studies that have looked at the genetic basis of ecotypic differentiation are often based on seedling transplantation (Lowry and Willis 2010; Fournier□Level et al. 2013) and so may exclude important aspects of divergence in earlier life stages (Donohue et al. 2010; Donohue 2014). This is surprising as early life stages are often crucial because this vulnerable period of development can suffer from high mortality rates (Donohue et al. 2010; Kitajima and Fenner 2000), thus selection may be severe during this part of the life cycle. Additionally, early life experiences often have non-linear effects on later life history traits (Beckerman et al. 2002). Thus, it may be difficult or impossible to fully understand adaptation if we ignore juvenile stages.

Seed size plays an important role on earlier growth and establishment. It connects the ecology of reproductive strategies with seed dormancy, seedling establishment and vegetative growth (Grime et al. 2014; Shipley et al. 1989; Leishman and Westoby 1992; Rees 1996). Thus, selection might favor different seed size strategies across heterogeneous habitats. For example, plants adapted to xeric habitats tend to have larger seeds compared to those occurring in mesic habitats (Baker 1972). A strategy producing fewer large seeds might ensure successful establishment by generating vigorous seedling growth, perhaps timed with pulses of seasonal precipitation. In contrast, a strategy producing many small seeds might arise in environments with highly predictable patterns of rainfall or where establishment and success are limited by other resources. Several studies have shown a positive correlation with seed mass and seedling root and shoot growth in different species (Jurado and Westoby 1992; Lloret et al. 1999; Wulff 1986). However, it is not clear whether seed and seedling traits jointly evolve by common pleiotropic effects or strong linkage disequilibrium or if these traits evolve independently.

Additionally, seed dormancy is another important trait tightly linked to plant life history (Venable 2007; Childs et al. 2010). A central idea is that dormancy is a bet-hedging strategy for reproduction in an unpredictable and heterogeneous environment (Venable 2007; Childs et al. 2010). In systems with strong dormancy, specific environmental conditions must occur to break dormancy but also to ensure germination after dormancy is broken (Baskin and Baskin 1998). Empirical studies showed inconsistent relationships between seed mass and germination traits. For example, studies conducted with *Medicago truncatula* and *Brassica oleracea* showed no genetic correlation between these two traits (Dias et al. 2011; Bettey et al. 2000). Dechaine et al. (2014) showed a positive genetic correlation between seed mass and germination trait in *Brassica rapa*. In tomato, no genetic correlation was found between seed size and germination rate when tested in greenhouse condition (Khan et al. 2012) but a negative genetic correlation was observed when tested under different nutritional conditions (Geshnizjani et al. 2020). Thus, it is not clear whether seed dormancy can evolve independently, or dormancy is tightly linked with seed mass variation in natural systems.

While seed and early life stages are likely critical in plant performance, our knowledge about the genetic basis of these, their plasticity, or their role in local adaptation is limited to only a few systems (Donohue et al. 2010; Donohue 2014; Clauss and Aarssen 1994). Seed mass and dormancy exhibit considerable phenotypic variation in nature (Bradford and Nonogaki 2008; Baker 1972; Harel et al. 2011; Lacerda et al. 2004; Bentsink et al. 2010). This variation might result from phenotypic plasticity or genetic differentiation. Based on habitat and seed size correlations, patterns in nature suggest seed size evolution is adaptive and is under stabilizing selection. These conclusions are mostly based on evidence from experiments with crop plants grown in controlled environments (Silvertown 1989). Studies with wild plants often show marked phenotypic plasticity and low heritability of seed size (Stanton 1984; Primack and Antonovics 1981). If the variation is not plastic and instead caused by genetic differentiation, then exploring genetic architecture underlying this natural variation would help understand the evolutionary forces driving seed size related phenotypic variation. It also helps to understand how adaptive evolution shaped genetic variability for these traits.

Here, we report on work exploring seed trait ecology and genetics in *Panicum hallii*, a self-fertilizing C4 perennial bunch grass species. *P. hallii* is distributed across the southwestern part of the US and northern Mexico (Smeins et al. 1976; Waller 1976; Hatch et al. 1999; Lovell et al. 2018b; Gould et al. 2018) with population genetic structure and putatively adaptive differentiation across its range (Lowry et al. 2013; Palacio-Mejía et al. 2021). However, the greatest differentiation within *P. hallii* occurs between ecotypes adapted to lowland/coastal and upland/southwestern regions (Lowry et al. 2015). These two ecotypes are

*P. hallii* var. *hallii* and *P. hallii* var. *filipes. hallii* is the more widespread ecotype and occurs primarily in xeric upland calcareous soils whereas *filipes* is located primarily in mesic and seasonal wet areas (Lovell et al. 2018a). In our study, we used field experiments to ask whether seed traits exhibit phenotypic plasticity in response to xeric and mesic habitat variation. Second, we used field seed addition experiments to test for local adaptation and tradeoffs based on seedling recruit and survivorship patterns. Finally, we used detailed growth chamber studies with a recombinant inbred population to explore the genetic architecture underlying ecotypic divergence in seed and seedling traits using quantitative trait loci (QTL) mapping. Together, our results provide insight in possible avenues of local adaptation mediated by seed traits.

## Methods

### Reciprocal transplantation

For plant material, we picked two representative inbred genotypes of the upland and lowland ecotypes (*hallii* and *filipes*, respectively) of *Panicum hallii* that have been used in earlier genome assembly efforts (Lovell et al. 2018b). We reciprocally transplanted them to their lowland (Nueces Delta Preserve; NDP, Odem Texas) and upland natural habitats (Brackenridge Field Lab; BFL, Austin, Texas) to evaluate seed trait plasticity. The Austin site represents a typical upland habitat with dryer and shallower soil, whereas the Odem site represents the lowland coastal prairie habitat. We germinated seedlings in one-inch pots in a greenhouse located in University of Texas, Austin and grew them to 20 days of age. We transferred 300 seedlings to the field sites in Fall 2018 (75 uplands + 75 lowlands = 150 × 2 sites = 300). After transplanting, we did not water or fertilize them but visited both sites in 30-day intervals until November 2018, when we collected seeds from both sites. We later randomly picked ten plants from each genotype to measure seed mass and dormancy. In total, we measured 40 seed lots (two genotype × two locations × ten replicates). Seed mass was calculated by weighing 100 seeds per line using an analytical balance (Mettler Toledo, Ohio, USA). We established a germination assay in a lab growth chamber with 15 seeds per petri dishes (unit of replication, total 40 petri dishes) and calculated germination percentage for these lines.

To study *P. hallii* recruitment, we conducted another reciprocal experiment at the same field sites. We established fourteen plots (0.5 × 0.5 m^2^) at each site haphazardly arrayed over native habitat. In February 2019 we added 100 seeds per plot from either *hallii* or *filipes* at each of the two field locations (two genotypes × seven replicates × two field sites = 28 plots). We visited each site every 15 days from the date of seed addition and collected two seasons (spring and fall 2019) of seedling and adult (reproductive) recruitment data. During each visit, plots were inspected for new seedlings and tagged using a mini rubber band. We also found other grass species emerging with *P. hallii* recruitment. To minimize confusion and misidentification, we also planted ten *hallii* and ten *filipes* in one-gallon pots prepared at both field sites. We used these pots as a reference while identifying seedlings in the field. Once we identified and marked seedlings, we followed them from early establishment to flowering. We used flowering as a metric to count adult plants.

### RIL population, linkage map and experimental setup

We used a recombinant inbred line (RIL) population generated from upland (*hallii;* HAL2 genotype) and lowland (*filipes;* FIL2 genotype) parental genotypes. A detailed description of the development of the RIL population and the genetic map construction are discussed in Khasanova et al. (2019). In brief, the RIL population was generated following a single seed descent method from F_2_ progeny of a single F_1_ hybrid to the F_7_ generation. A total of 335 RILs were re-sequenced up to 30x coverage and used to build a high-density linkage map after mapping against the *filipes* reference genome. The final map consists of 722 markers in 9 chromosomes. For this study, 295 RILs and both parental lines were collected from well-watered plants grown as a seed increase at the Brackenridge Field Laboratory, Austin, Texas in September 2016. Seeds were collected from three maternal parents from each RIL. We carefully separated good seed from chaff, detritus, or undeveloped seeds and kept them at room temperature for 6 months before starting the experiment trials.

We utilized a large walk-in growth chamber experiment to control environmental conditions throughout the experimental trials. Growth chamber temperature was set at 28°C during light and 22°C during dark conditions. The photoperiod for the chamber was set at 12 hrs light/12 hrs dark. Seeds were germinated in petri dishes (25×100 mm) with sterilized sand. We used 60 g of sands for each petri dish and added 12 ml tap water to each dish. Water was sprayed on the upper lid of the petri dishes to keep the inner environment moist, and each dish was sealed with parafilm. The experiment was replicated across three temporal blocks in which each block contained a single petri dish (our unit of replication) of each genetic line. In each petri dish, 15 seeds per lines were used. The complete experimental design was as follows: 295 RILs + six petri dishes of *hallii* + six petri dishes of *filipes* × three replication = 921 petri dishes.

Because we consistently observe strong seed dormancy in the *hallii* ecotype, we wanted to make sure an inference of dormancy in *hallii* is not simply the result of inviable/sterile seeds. Here, we applied scarification and found it to be effective for breaking dormancy in the upland *hallii* ecotype. We examined all non-germinated seeds from this germination assay to evaluate viability and dormancy by forcing germination after removing seed coats. We removed the seed coat by gently using sandpaper (P100) and then germinated them in petri dishes.

### Phenotypic traits measurement at the seed, germination, and earlier growth stage

We measured six seed size and earlier seedling growth traits from our growth chamber experiment. Seed mass (SM) was measured by taking 100 seed weight per RIL line using an analytical balance. We recorded the day of first germination (GT) and germination percentage (GP) for each RIL as assayed in petri dish trials. Viability experiments with some random RIL lines and parental lines confirm that un-germinated seeds were dormant, and as such we view GP as a proxy for the degree of seed dormancy. Shoot length was measured twice on a representative seedling in a dish, at 5 days (SL5D) and at 10 days (SL10D) relative to their GT. Root length was also measured on a representative seedling per petri dish at 10 days (RL10D) relative to GT. A detailed description of our phenotypic data measurement is in supplementary file S1.

### Data processing, statistics, and QTL mapping

We took two approaches to analyze our reciprocal field experiment data. To analyze seed mass and germination percentage in the first reciprocal transplant experiment, we fit linear models with genotype (G), site (E) and their interaction (GxE) as the effect variables. To analyze counts of seedling and adult recruitments in the second reciprocal transplant experiment, we fit generalized linear models with a Poisson distribution and log link. These models took the same form as the linear model above. We determined that the Poisson GLM adequately fit the data using the Pearson goodness of fit statistics. All models were fit in JMP version 15.1.0 (SAS Institute, Cary, NC).

For quantitative genetic studies, we fit linear models for the measured phenotypic traits treating RIL genotype as a fixed effect. We also included temporal block in the model as a fixed effect covariate when the block had a significant effect on phenotypic traits. Most of our collected phenotypes except SM and SL10D had a significant block effect. We used RIL means as estimates of breeding values to calculate the bivariate genetic correlation between traits using ‘multivariate’ function in JMP version 15.1.0. Broad-sense heritability was estimated using h2boot software fitting one-way ANOVA among inbred lines with 1000 bootstrap runs (Phillips and Arnold 1999). Parental trait divergence was estimated by mean differences and significance tested with a *t*-test in SAS. Resulting p-values were adjusted for multiple testing via the Benjamin and Hochberg’s (1995) method of False Discovery Rate (FDR) estimation.

Fraser (2020) noted that the pattern of trait variance in recombinant populations can provide insight into whether the parental trait divergence is consistent with neutral divergence or more likely the result of natural selection. The approach is a generalization of the QTL sign test (Orr 1998) and rests on the realization that the phenotypic distribution in a segregating/recombinant population can be treated as a null model for the phenotypes expected under neutral evolution. In this framework, the pattern of underlying QTL effects under strong directional selection is expected to be complementary (in a consistent direction) if the trait has experienced consistent strong directional selection leading to parental divergence, whereas a mixed of positive and negative effects is expected if the trait evolved by neutral processes. By extension, if a trait has experienced strong directional selection driving parental divergence the variance among parental lines is expected to be larger than the segregating variance in the recombinant population, The v-test was performed as described in equation 2 of Fraser (2020) for a RIL. In brief, to calculate *v*, we first estimated the trait variances within and between parents of the cross. Then we estimated the variance among RIL means and used a *c* value of 1 (Fraser 2020).

All measured morphological traits in the RIL population were normally distributed except for SL10D and RL10D. These traits were log transformed before QTL mapping. We performed QTL mapping on RIL breeding values by stepwise fitting of multiple QTL models in R/qtl package (Broman et al. 2003). The input file for QTL mapping including the linkage map and phenotypic traits is in supplementary file S2. We used two functions to determine QTL positions and for the estimation of additive and pairwise epistatic effects. We first calculated penalties for main effects and their interactions using Haley-Knott regression by 1000 permutations of the scan-two function for each phenotypic trait. We then performed a forward/backward stepwise search for models with a maximum of six QTL that optimized the penalized LOD score criterion using the stepwise procedure. Significance thresholds were determined at an experiment wide Type 1 error rate of α = 0.05. Confidence intervals for significant QTLs were calculated as the 1.5 LOD drop interval by using the *qtlStats* function from *qtlTools* (github.com/jtlovell/qtlTools).

## Results

### Seed mass and earlier growth traits varied significantly between parental lines

Parental lines exhibited considerable phenotypic divergence in seed size, dormancy, and earlier growth traits (Fig. 1). In growth chamber experiments, parental lines significantly differed for all six measured traits (Table 1). The upland ecotype had higher values in all traits compared to the lowland ecotype except for the GP. For example, *hallii* had 56% greater SM than *filipes* and the GP in *filipes* was 63% greater than *hallii*. The mean difference in GT between ecotypes was 60 hrs (± 2.7), with *filipes* germinating earlier than *hallii*. Early shoot and root length in *hallii* were greater than *filipes*. Shoot length after 5 days was 10% greater in *hallii* and the differences increased to 24% at the 10-day measure. Ten days old root growth was 73% greater in *hallii* (Table 1, supplementary file S3).

**Table 1:**
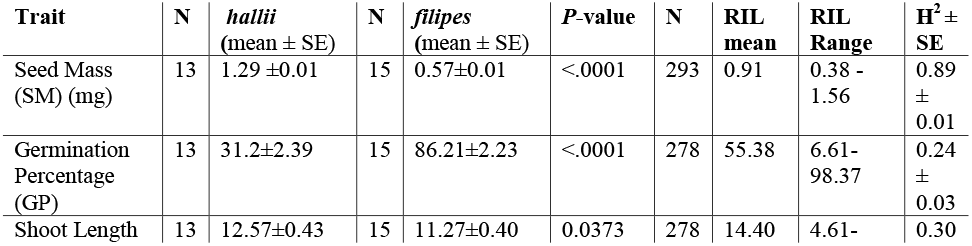
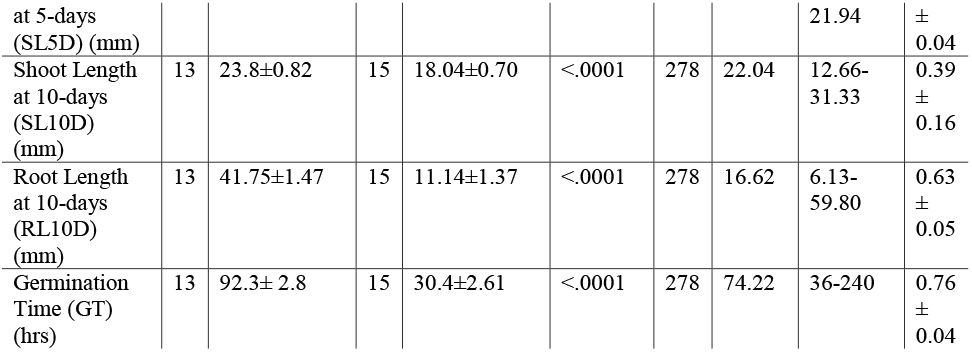
Descriptive statistics for phenotypic traits for *hallii* and *filipes* parental lines and the recombinant inbred population (SE= standard error).

**Fig. 1:**
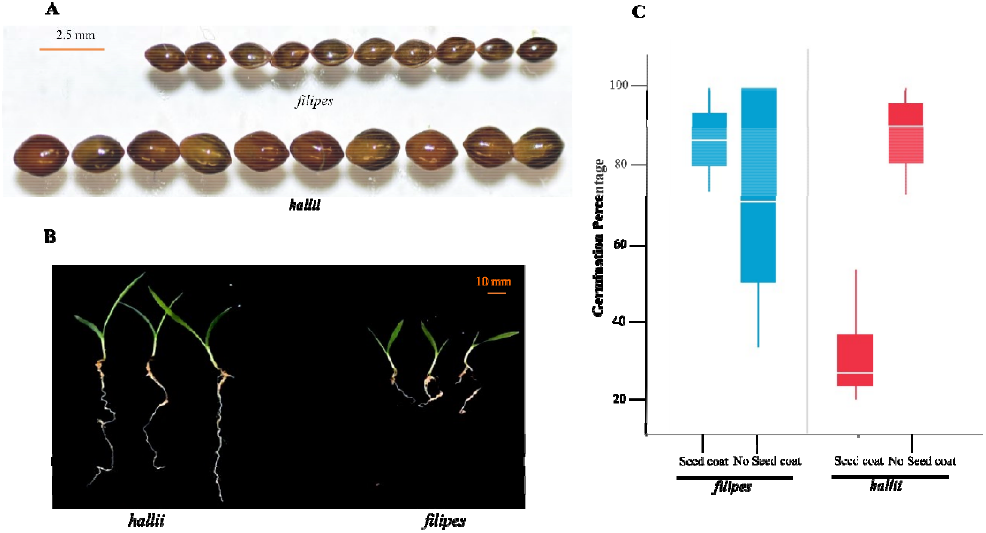
Seed and seedling size differences and seed viability tests between *P. hallii* lowland and upland ecotypes. 1A shows the representative differences in seed size between parental lines. This image was taken with 10 random seeds for both parental lines. 1B shows seedling growth differences for these parental lines at 10 days from the day of first germination. The upland ecotype grew larger than the lowland ecotype in both belowground and above ground biomass. 1C shows the result of seed viability test between *hallii* (red) and *filipes* (blue). Ungerminated seeds were tested for germination by removing seed coats. The Y-axis corresponds to the percent of germinated seeds and the X-axis provides the genotype information. Upland *hallii* has a higher germination rate compared to *filipes* if seed coats are removed.

Additionally, we tested germination percentage for the non-germinated seeds by removing seed coats to check if the observed dormancy is due to seed viability problem. Our data showed that a scarification technique effectively increased seed germination for the dormant *hallii* seeds. Upland *hallii* seeds germination was increased to 88.2% after removing seed coats, a 57% increase compared to their regular germination in the absence of seed coat removal (Fig. 1, supplementary file S4). This result confirmed that the apparent seed dormancy in the upland ecotype was not due to seed viability issues and may be related to inhibition associated with the seed coat.

### Seed plasticity, seedling recruitment and adult establishment in parental lines

To explore the impact of seed development in native habitats on seed traits, we collected seeds from field plantings at reciprocal sites. We tested for genotype (G), site (E) and genotype-by-site (GxE) interaction effects on seed mass and germination traits. In this framework, we interpret significant G as evidence for ecotype divergence in seed traits, significant E as habitat driven seed plasticity, and GxE as ecotype variation in plastic responses. Seed mass revealed significant genotype main effects (*P* < 0.0001) but did not show significant effects of reciprocal sites (*P* = 0.86) or GxE interaction (*P* = 0.27). We used field collected seed in lab germination experiments to evaluate impacts on seed dormancy. As with seed mass, we observed strong genotype difference in germination percentage (*P* < 0.001), but no significant differences between reciprocal sites (*P* = 0.38) or for GxE interaction (*P* = 0.72) (Fig. 2, supplementary file S5). These results indicate minimal phenotypic plasticity but strong genetic divergence in seed size and germination characteristics.

**Fig. 2:**
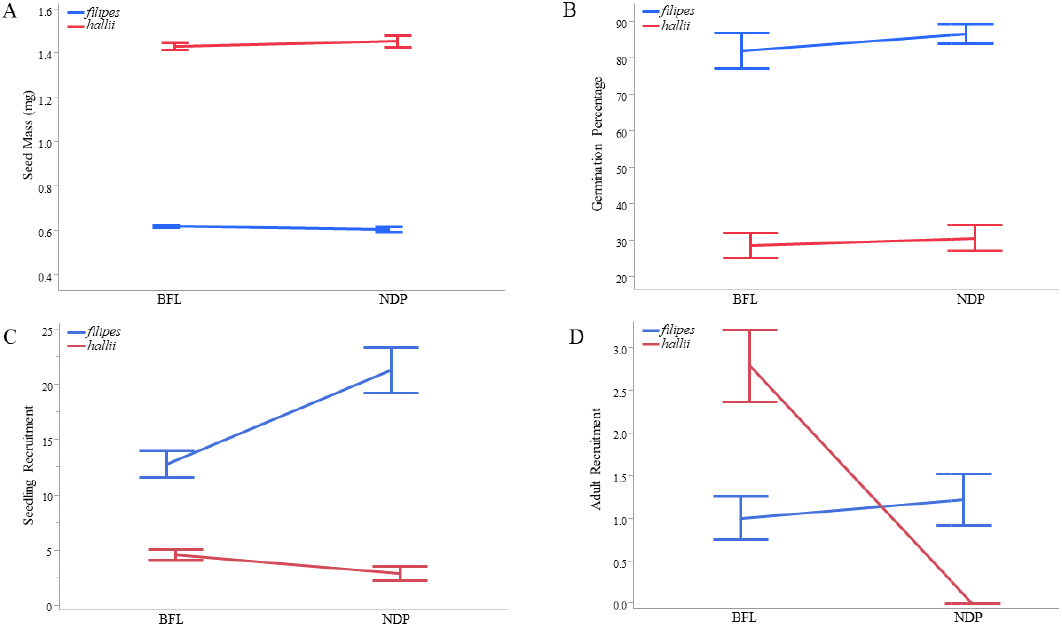
The effect of reciprocal transplantation on seed size, germination percentage, seedling recruitment and survivorship between the upland, *hallii* (red), and the lowland, *filipes* (blue), ecotypes. 2A-B show that the effect of reciprocal transplantation on seed size and germination percentage between parental lines. The differences between ecotypes were significant but the location does not affect seed mass or germination. 2C shows seedling recruitment and 2D shows the number of adult plants per plot in reciprocal habitats. The Y-axis corresponds the trait values and X-axis provides the location of the experiment (Brackenridge Field Lab, BFL, upland habitat; Nueces Delta Preserve (NDP, lowland habitat). Genotypes are indicated by color with *hallii* (red) and *filipes* (blue).

We also used seed addition plots to test for ecotypic differences in seed germination and seedling establishment. Overall, we observed many more seedlings (fivefold increase) from plots sown with *filipes* seed across both locations. Most recruitment occurred in the spring following the establishment of the experiment, with relatively few new seedlings in the following fall. Recruitment to the adult stage was low, especially for parental seed sown in foreign sites. We observed no *hallii* adult recruitment at the NDP (lowland) site and little *filipes* adult recruitment at either site. The majority of recruitment occurred for *hallii* seeds planted at the BFL (upland) field site. In factorial models, we detected significant GxE interaction for both seedling (*P* < 0.0001) and adult recruitment stages (*P* < 0.0001) (supplementary file S6). The form of the GxE suggests tradeoffs in early seedling recruitment favoring *filipe*s across habitats, but later adult establishment favoring *hallii* at the upland habitat. Together, these tradeoffs are consistent with a model of local adaptation driven in part by early life history stages. Our field experiments provide compelling evidence for strong genetic differentiation in seed traits and support for local adaptation through differential recruitment and survival in home versus foreign habitats.

### Heritability and genetic correlations among RILs

We estimated broad-sense heritability (H^2^) among RILs in the population as the proportion of total phenotypic variation due to genetic variation among RIL lines. All measured traits were heritable with heritability ranging from 24 to 89% (bootstrap-based significance, *P* < 0.001). GP was the least (0.24) and the SM was the highest heritable traits (0.89). The heritability of GT and RL10D were also notably high, 0.76 and 0.63 respectively. The estimated heritability for SL5D and SL10D were 0.30 and 0.39 respectively (Table 1).

We observed significant genetic correlation between most of the phenotypes measured in the RIL population. SM was significantly positively correlated to SL5D (r_g_ = 0.15, *P* = 0.020), SL10D (r_g_ = 0.23, *P* = 0.001) and RL10D (r_g_ = 0.29, *P* = 0.001), but negatively correlated with GP (r_g_ = −0.44, *P* < 0.001). GT was negatively covaried with GP (r_g_ = −0.40, *P* < 0.001), SL5D (r_g_ = −0.36, *P* < 0.001) and SL10D (r_g_ = −0.21, *P* < 0.001). GP was positively correlated with SL5D (r_g_ = 0.34, *P* < 0.001) and SL10D (r_g_ = 0.17, *P* = 0.009) but negatively correlated with RL10D (r_g_ = −0.13, *P* = 0.035). Both shoot length traits (SL5D and SL10D) were also positively correlated (r_g_ = 0.67, *P* <0.001) among RIL. In general, RIL with larger seeds initiated robust shoot and root growth but fewer of their seeds germinated over the course of petri dish trials (Fig. 3).

**Fig. 3:**
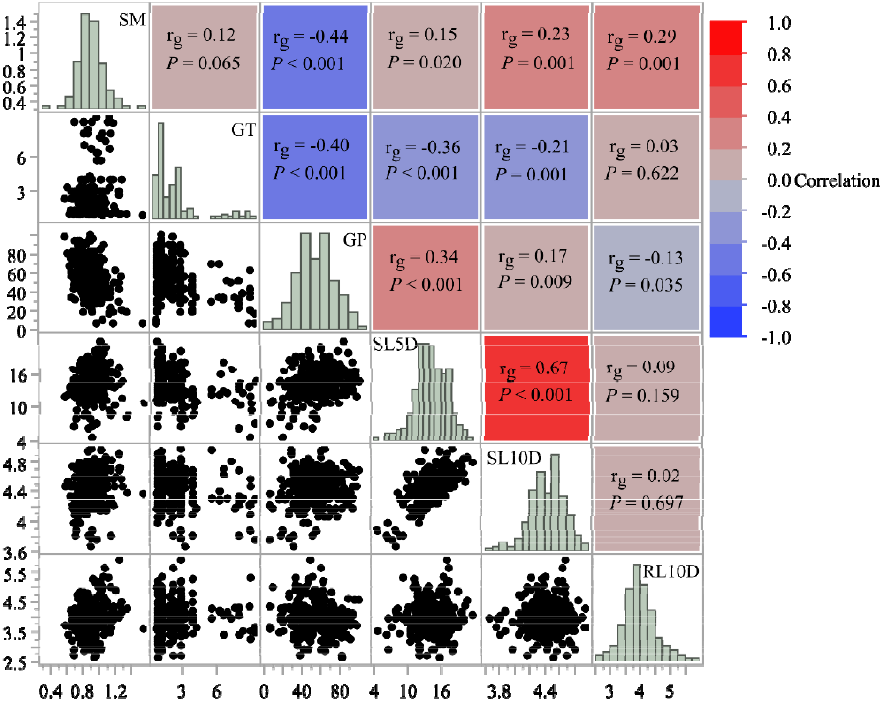
Genetic correlation coefficients (*r*_g_) and their pairwise significance (*p*-value) for seed mass, germination rate, shoot length and root length traits with RIL means. Seed mass showed an antagonistic relationship with germination percentage and showed a positive relationship with both shoot length and root length traits.

The Fraser *v*-test statistics are highly significant (*P <0*.*01*) for all traits except for SL5D (*P* = 0.57). Overall, the *v*-test framework identified phenotypic patterns for all six traits which are suggestive of directional selection underlying seed and seedling trait divergence among *P. hallii* ecotypes. In all cases, variances of parental trait values were greater than the variances among RILs (supplementary file S7). The most significant traits for directional selection were SM and RL10D (*P* < 0.0001), followed by GP (*P* = 0.0003), GT (*P* = 0.0018) and SL10D (*P* = 0.016). These results are consistent with strong directional selection pressures underlying the differences among these representative parents.

### Genetic architecture underlying seed size, germination, and seedling growth traits

We identified 20 QTL, including one epistatic QTL pair, from all measured traits using stepwise QTL models. SM, GP and SL5D comprised over half (75%, 15 QTL) of the identified QTL. Most of the QTL (13 out of 20 QTL) had overlapping confidence intervals with a QTL for at least one other trait, suggesting that in many cases phenotypic correlations are reflective of underlying genetic correlations resulting from physical linkage or pleiotropy. About 84% of the identified QTL affected phenotypes in the expected direction based on observed evolutionary divergences. For example, in all cases the *hallii* allele increased trait values relative to *filipes* alleles for SM, SL10D and RL10D. On the other hand, the *filipes* allele increased trait values for GP. We observed mixed QTL effects only for SL5D, where three of the QTL effects were increased by the *hallii* allele and two were increased by the *filipes* allele. Here, the direction of effects was more complex than expected based on ecotype divergence (Fig. 4). Details about individual QTL, including their location and confidence intervals, effects, and PEV can be found in supplementary file S8.

**Fig. 4:**
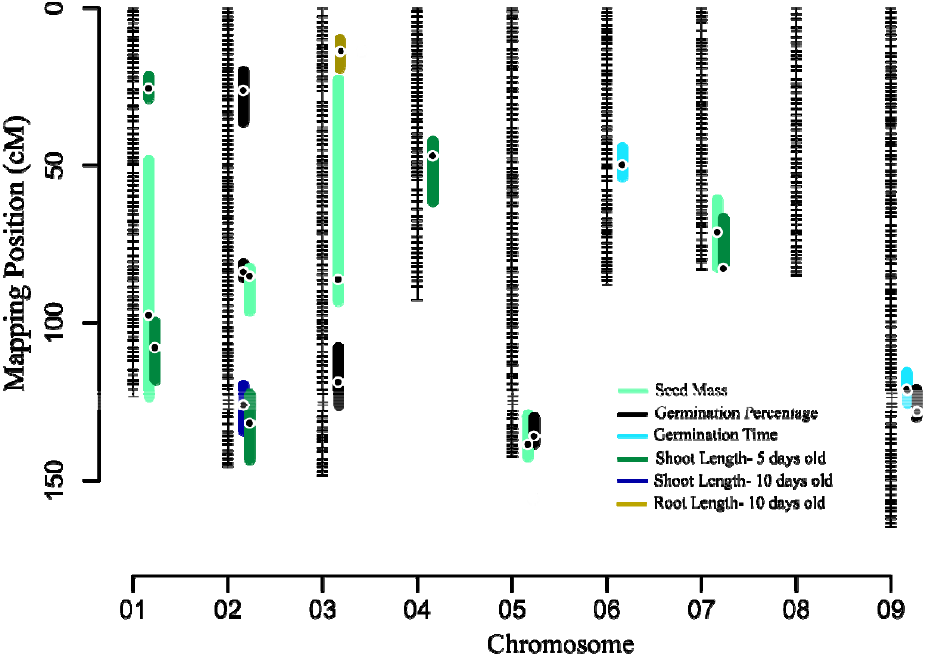
The RIL genetic linkage map of *Panicum hallii* generated from *hallii* × *filipes* mapping population. All identified QTL are presented as bars to the right of each linkage group. Each colored bar represents a QTL with length representing the 1.5-LOD drop confidence intervals. Black dots in the colored bars indicate the position of the QTL based on the maximum LOD score. GP = Germination Percentage, RL10D = Root length at 10 days, SA = Seed Area, SL= Seed Length, SL10D = Shoot Length at 10 days, SL5D = Shoot Length at 5 days, SM = Seed Mass, SR = Seed Roundness, SRR = Shoot Root Ratio at 10 days, SW = Seed Width and VI = Vigor Index.

Only single QTL were identified for SL10D and RL10D. The SL10D QTL overlapped with SL5D at Chr2@126 and the RL10D QTL had a unique position at Chr3@11.7. Despite these traits having relatively high heritability (Table 1), we only detected single QTL with a relatively low percent of variance explained for these traits (PVE < 10%), suggesting there may be many undetected loci controlling these traits or that our heritability estimates are inflated by maternal environmental or epigenetic effects.

### QTL colocalization, allelic effects, genetic trade-offs, and epistasis

Overall, we mapped 13 overlapping QTL regions. Colocalization of QTL could be indicative of loci with pleiotropic effects or genes in tight linkage that affect unique traits. We identified interesting and potentially functionally related combinations of QTL colocalization. For example, GP and GT trait QTL colocalized at Chr9@128 with antagonistic effects in the same direction as parental divergence. Here, the *filipes* allele increased trait values for GP but decreased values for GT. In another case, SM and GP shared two colocalized regions, one was at <u>Chr2@85.3</u> and the other one was at Chr5@130. In both cases the *hallii* allele increased trait values for SM and decreased GP trait values as expected from the observed evolutionary divergence between parental lines. We also observed QTL colocalization with mixed allelic recombination for same trait combination. SM and SL5D colocalized at two chromosomal regions. In the first colocalized regions at Chr1@108, *hallii* allele increased trait values for SM and decreased trait values for SL5D. But in the second region at Chr7@70, *hallii* allele increased traits values for both traits. We also found some expected traits colocalization. For example, shoot length measured at two time points, SL10D and SL5D, colocalized at Chr2@126 having additive effects in the same direction of parental divergence where the *hallii* allele increased trait values for both traits (supplementary file S8).

Only 25% of identified QTL (5 out of 20) had unique position in the linkage map. From the trait measured, SL5D had two unique QTL (1 @ 25.7 and 4 @ 47.1). GP and GT had a single unique QTL occurring at 2 @ 26.3 and 6 @ 50 respectively (Fig. 4, supplementary file S8).

Finally, we detected one pairwise epistatic interaction for GT occurring between QTL on Chr6 and Chr9 (Fig. 5). Here, individuals that are homozygous for the *hallii* allele at both loci have delayed germination, but this pattern is masked in any of the alternative hybrid combinations.

**Fig. 5:**
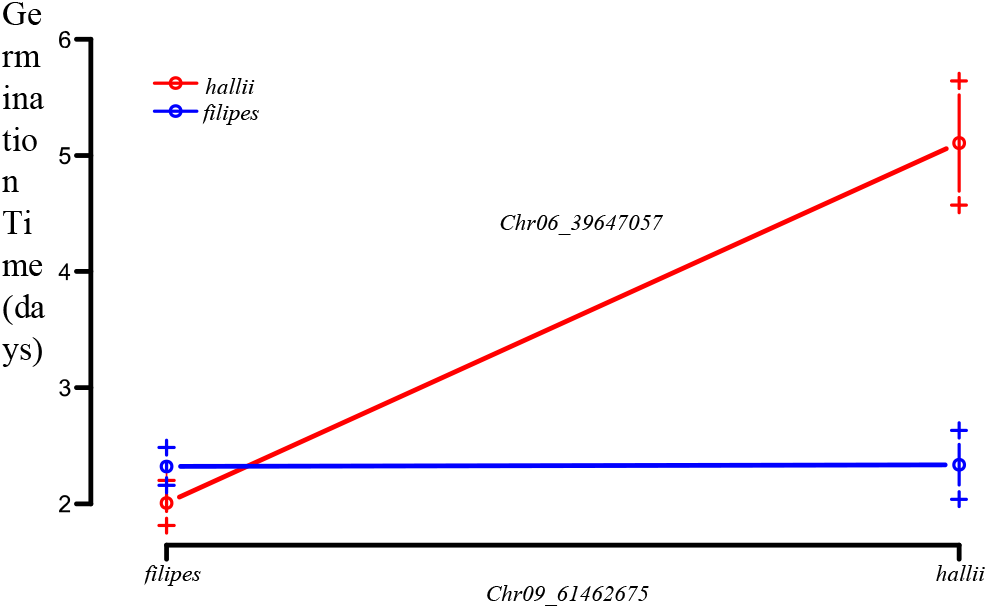
Pairwise epistatic QTL for the day of first germination (GT) between Chromosome 6 and 9 in the *P*.*hallii* RIL population. Plotted points indicate two-locus genotype means ± 1SE for the two loci containing GT between Chr6 and Chr9.

## Discussion

Populations adapting to heterogeneous environments will develop genetic differences over time in response to selection. This adaptation may ultimately lead to ecotype formation with distinct morphological and/or physiological characteristics that provide an environment-specific fitness advantage. In this study, we examined ecotypic divergence between xeric and mesic ecotypes at the seed and seedling stages in *P. hallii* to evaluate early life stages that may contribute to local adaptation. Field experiments reveal little plasticity but support a hypothesis of local adaptation driven by tradeoffs in seedling recruitment and adult establishment. We mapped QTL for six traits with several large effect QTL colocalizing to a relatively small number of genomic regions. Overall, the direction of QTL effects is predominantly in the direction of putative adaptive divergence. This pattern is consistent with a hypothesis of strong directional selection on seed related traits (Rieseberg et al. 2002; Orr 1998) as indicated by significant *v*-tests (Fraser 2020). We also identified clear genetic trade-offs between seed mass and germination percentage at two QTL based on QTL localization. These data suggest that seed size, dormancy and seedling vigor traits may have jointly evolved in the process of ecotype divergence.

### Parental divergence and trait plasticity

Plant populations are generally limited by either the abundance and quality of seed for seedling recruitment or by the availability of microsites for seedling establishment. One hypothesis is that aspects of habitat quality and the local competitive environments may therefore play key selective roles in the evolution of seed and seedling phenotypes. Considering this theory, it is interesting to speculate on the patterns of ecotypic divergence in *P. hallii. P. hallii var. hallii* is a widespread ecotype and occurs primarily in xeric habitats in rocky, dry, and calcareous soils (Smeins et al. 1976; Waller 1976; Hatch et al. 1999) and with relatively low ground cover or competition. In contrast *var. filipes* is located primarily in mesic and seasonal wet areas (Gould 1975; Waller 1976) in dense coastal prairie habitats. Our data show that *hallii* produces larger seeds with delayed germination and strong dormancy compared to *filipes*. In addition, *hallii* roots grew 73% longer than *var. filipes* roots after ten days of seedling growth (Fig. 1, Table 1). Typically, increased root growth from large seeds would result in an establishment benefit in resource poor sites. Thus, having larger seed mass, high dormancy, delayed germination time and faster root growth may be selected for *hallii* to survive hazards of establishment in harsh southwestern xeric habitats. Reciprocal plantings of ecotypes revealed that local conditions have little impact on seed mass or germination percentage. Our seed addition experiment indicates that the lowland ecotype (*filipes*) is most likely limited by seed or microsite availability in general, while the upland ecotype (*hallii*) exhibited more striking habitat dependent establishment (Fig. 2, supplementary file S5, supplementary file S6).

Our results support the model developed for maintenance of seed size diversity by Muller-Landau (2010), where species coexist in heterogenous habitats by a tolerance-fecundity trade-off. Under this mechanism, species with larger seeds win in a stressful environment (e.g., too dry, too shady etc.) due to their higher tolerance to stress while small seeded species win in competitive habitats due to their higher fecundity. We find *filipes* commonly in competitive coastal prairie environments with a high density of other grasses (e.g., *Bouteloua rigidiseta, Nassella leucotricha*). The upland strategy centers on producing fewer larger seeds with high dormancy, and the lowland strategy centers on producing many small seeds with low dormancy. Larger seed mass in the upland ecotype is probably under strong selection to improve establishment in dry and calcareous soil and seed number is possibly under strong selection in the lowland ecotype as a mechanism to provide opportunities for establishment. Here, selection may have favored the production of a large number of small seeds with attributes allowing establishment in rare disturbed patches of habitat. Future field studies exploring the impact of competitive vegetation, disturbance, or seasonality on seedling establishment will help better reveal possible mechanisms of selection on seed traits in *P. hallii*.

### The genetics of ecotypic divergence

Ecotypic divergence caused by adaptation to xeric and mesic habitats provides ample evidence of ecological functions that contribute to the process of speciation (Clausen 1951; Kruckeberg 1986; Rajakaruna 2004; Lowry 2012; Lowry et al. 2015). Understanding the underlying genetic basis of ecotypic divergence helps to better understand the degree to which adaptive evolution was constrained or facilitated by the structure of genetic variation. In this study, we found colocalization of QTL for a common set of traits involved in xeric and mesic ecotypic divergence, suggesting that these traits may jointly evolve. We observed that seed mass QTL colocalized with all other measured traits, suggesting that in many cases seed mass, seed investment and earlier growth-related traits may function as an integrated suite of functional traits.

We also found evidence for genetic trade-off between seed-based life history traits. Most of the discovered QTL clusters in this study had antagonistic effects across traits. The finding of antagonistic additive effects for QTL clusters implies that seed based related traits are under strong selection between

*P. hallii* lowland and upland ecotypes. For example, seed mass and germination percentage colocalized at two chromosomal regions (Chromosomes 2 and 5) where the *hallii* allele increased seed mass but decreased germination percentage. Our result suggests that *P. hallii* ecotypes may have experienced strong selection pressures to evolve either large seed/strong dormancy or small seed/weak dormancy trait combinations. Additionally, GP and GT colocalized at Chr9 where the *hallii* allele decreased GP but increased GT. The observed genetic covariation in the RIL population suggests that the major axis of standing genetic variation would support the rapid evolution of these trait combinations, while potentially constraining evolution of the opposite combination of traits. In addition, large seeds with vigorous root growth might be favored in dryer and resource limited habitats (Fig. 3). The Fraser *v*-test also supports this inference; the highly significant *P*-values are consistent with directional selection for these trait combinations while adapted to lowland and upland habitats. Thus, our mapping data support the idea that large seeds/ dormancy and longer root growth trait combination may have been an important adaptation for *hallii* as it diverged from a coastal ancestor and invaded dry, calcareous habitats across the southwest.

We have consistently found that upland and lowland ecotype differentiating traits colocalized in *P. hallii* to a relatively small number of chromosomal regions. Lowry et al. (2015) mapped five QTL to a locus on chromosome 5 with traits that are involved in ecotypic divergence using an earlier F_2_ population. These traits are mainly associated with flowering time, tiller characters and leaf width. In our study, two traits (seed mass and germination percentage) colocalized to this region. Interestingly, all five traits that Lowry et al. (2015) mapped had additive effects in the same direction where the *filipes* allele increased the trait values. Khasanova et al. (2019) also mapped five shoot and root traits to the same region. In that study, all traits also had additive effects in the same direction with the *filipes* allele increasing trait values. In our study this QTL had contrasting effects on seed size and germination, in that the *hallii* allele increased seed mass trait values and decreased germination percentage. Our study therefore provides further support for a major pleiotropic region that contributes to ecotype differentiation in *P. hallii*.

## Conclusions

Seed biology plays an important role in plant ecology. It tells us how plant species invest resources into reproduction, and how natural selection has given rise to an incredible diversity of reproductive strategies. In this study, we explored several facets of ecotype differentiating traits from a seed biology perspective. Our study reinforces a pattern that have been observed in multiple species, wherein genetic studies find that ecotype differentiating traits colocalize to common genomic regions (Latta and Gardner 2009; Lovell et al. 2013; Lowry et al. 2015; Khasanova et al. 2019; Hall et al. 2006; Lowry and Willis 2010). Once fine mapping or transformation becomes feasible in this species, further work can identify candidate genes to gain better insight into the molecular details underlying local adaptation and ecotypic divergence in *P. hallii*. Overall, our data support the functional integration of seed size and seed dormancy traits, an exciting next step will be to evaluate the fitness and population impact of the seed variation in different habitats.

## Acknowledgements

Thanks to Brandon Campitelli, Allison Hutt & Shayan Noshir Bhathena for helping during seed collection. Taslima Haque and Juan Diego Palacio Mejia helped with the initial set up of the experiment. Robert Heckman gave important comments and valuable insights during the work, analyses and manuscript writing. Alice MacQueen provided feedback on the manuscript. Thanks to Xiaoyu Weng, Bhaskara Badiger and Joseph Edward for helpful suggestions. Thanks to Jake Herring, Director of Land Conservation, from Coastal Bend Bays and Estuaries Program for allowing us to conduct field plantings at the Nueces Delta Preserve, Odem Texas. Rob Plowes provided space at the Brackenridge Field Lab site at UT Austin. Shane Merrell provided technical support while setting up the experiment in the greenhouse facility. This research was supported by an NSF Plant Genome Research Program Grant (IOS-0922457) to TEJ. Authors also acknowledge Texas Ecolab support to SR for conducting field work.

## Author contributions

SR and TEJ designed the experiment. SR conducted the experiment, analyzed the data and wrote the manuscript. TEJ provided feedback on analyses and edited the manuscript.

